# Digital Imaging and Vision Analysis in Science Project improves the self-efficacy and skill of undergraduate students in computational work

**DOI:** 10.1101/2020.10.26.353987

**Authors:** Tessa Durham Brooks, Raychelle Burks, Erin Doyle, Mark Meysenburg, Tim Frey

**Author notes:** Corresponding author (TDB).

## Abstract

In many areas of science, the ability to use computers to process, analyze, and visualize large data sets has become essential. The mismatch between the ability to generate large data sets and the computing skill to analyze them is arguably the most striking within the life sciences. The Digital Image and Vision Applications in Science (DIVAS) project describes a scaffolded series of interventions implemented over the span of a year to build the coding and computing skill of undergraduate students majoring primarily in the natural sciences. The program is designed as a community of practice, providing support within a network of learners. The program focus, images as data, provides a compelling ‘hook’ for participating scholars. Scholars begin the program with a one-credit spring semester seminar where they are exposed to image analysis. The program continues in the summer with a one-week, intensive Python and image processing workshop. From there, scholars tackle image analysis problems using a pair programming approach and finish the summer with independent research. Finally, scholars participate in a follow-up seminar the following spring and help onramp the next cohort of incoming scholars. We observed promising growth in participant self-efficacy in computing that was maintained throughout the project as well as significant growth in key computational skills. DIVAS program funding was able to support seventeen DIVAS over three years, with 76% of DIVAS scholars identifying as women and 14% of scholars being members of an underrepresented minority group. Most scholars (82%) entered the program as freshmen, with 89% of DIVAS scholars retained for the duration of the program and 100% of scholars remaining a STEM major one year after completing the program. The outcomes of the DIVAS project support the efficacy of building computational skill through repeated exposure of scholars to relevant applications over an extended period within a community of practice.

## Introduction

Science, technology, engineering, and mathematics (STEM) professions, even those not traditionally steeped in quantitative models and data analysis, increasingly require computational competence [1]. In particular, the natural sciences have experienced significant increases in the amount of data generated by increased computing power, cheaper and more rapid sequencing technologies, and the rise of interdisciplinary fields such as personalized medicine, phenomics, digital agriculture, and climate science. Computation has become so ubiquitous and necessary across the natural and physical sciences that it has been referred to as the “third pillar of the scientific method,” along with theory and experimentation [2]. A career in the natural sciences increasingly requires that professionals are comfortable with basic computational skills and quantitative analysis [3–5]. Beyond this, modern scientific exploration may require the design of new software by developers with both specific content knowledge and computational skills. As a potential “end user”, a biologist, chemist, physicist, etc. has the content knowledge, but may need computational skills training [6,7]. Across the broad range of STEM disciplines, too few students are being trained in computational and quantitative skills that would enable them to develop useful software. In particular, undergraduate students in the life sciences may be resistant to developing quantitative or computational skills due to previous negative experiences or a perception that they “aren’t good at” mathematics or computers [8]. The result of these factors is a mismatch between the skills needed for success in research or industry positions and the skills possessed by graduates and young professionals starting these positions.

To address this mismatch, we conceived of the Digital Imaging and Vision Applications in Science (DIVAS) Project. This year-long program was designed as a guided ‘onramp’ to develop computational skills within a community of practice that would contribute to participants’ STEM career success. The overall goal of the DIVAS project is to develop, utilize, and test interventions that will engage and train STEM undergraduate students in computing - especially students that do not traditionally participate in computer science curriculum. DIVAS interventions present students with visually-appealing image-based problems relevant to the disciplines they are majoring in, thereby making the skills we aim to develop eminently practical. Importantly, it is relatively easy to capture images with high spatial, temporal, and spectral resolution, with images being increasingly used as data in scientific, clinical, and engineering settings [9–12]. While images are relatively easy to obtain, extracting useful information from them commonly presents technical barriers that lead to processing bottlenecks. Although the collection of large datasets has become rather commonplace, scientists of various career stages may lack the computational skills to analyze these data independently or may have limited access to productive collaborations with computer scientists or other specialists. Early introduction to computational approaches, along with frequent practice, enables a person new to computing to take advantage of training resources to develop critical skills and to form effective collaborations [13–15]. Studies of computer science courses that present instructional concepts in the context of digital images, videos, or music - i.e. “media computation” [16] - improves retention of women and non-computer science majors in these courses [13,15,17,18].

Just as computation-in-context supports student gains, so do communities of practice and learning communities. Both types of communities, which can be quite distinct depending on their specific model [19], are often used interchangeably to describe a community for sharing, developing, and/or maintaining knowledge, skills, and practices within which membership ranges from novices to seasoned experts. For students, participation in such communities has been shown to boost academic performance, self-efficacy, sense of belonging, STEM identity, retention, and graduation rates [20–23]. In the DIVAS Project, cohorts of novices work side-by-side with faculty mentors, and their more experienced student peers, to themselves become more advanced practitioners via legitimate peripheral participation [24]. Importantly, the DIVAS Project models the reality of the modern computational work environment, which is soundly a team-based endeavor. This counters the stereotype that such work is largely solitary.

The general hypothesis of the DIVAS Project is that gradual, scaffolded exposure to - and practice with - computational tools, centered on accessible and relevant applications, and implemented in both simulated and authentic supportive professional environments, will impact student self-efficacy, computational competency, and career path interest. We have taken the approach of emphasizing growth in self-efficacy toward computing as the first necessary indicator of growth in computational skill [25–27]. We also posit that as participants become more familiar with computational tools, they will additionally show more interest in career paths that would utilize said tools. Though our pilot program was restricted in size, its positive impact on participants suggests that DIVAS program elements are well-suited to our broader goals of fostering computation skills within a community of practice. We describe our approach here both as a guide and an invitation. We hope to form new DIVAS partnerships to broaden the DIVAS community and enable additional study on the efficacy of the approach we have taken.

## DIVAS Program Elements

To explore our hypothesis, a pathway of interventions was designed that comprise our programmatic ‘onramp’ (Fig 1). Each cohort of DIVAS scholars was introduced to our community of practice via a one-credit, spring semester seminar (DIVAS Seminar I) and engagement with the DIVAS Slack team. Work continued in the summer with a week-long coding workshop, followed by a four-week long paired-programming session that allows DIVAS scholars to put their recently acquired skills to use. DIVAS Scholars can participate in an additional three weeks of research with DIVAS faculty to conclude their summer activities. In the fall semester, DIVAS Scholars returning to research - or starting new computing projects - continue engaging with community members using our Slack team. During the following spring, the cohort takes DIVAS Seminar II. As with other team science endeavors, the DIVAS community is offline and online, with Slack and Zoom playing significant roles in communication, project management, co-working sessions, team meetings, etc. In the sections that follow, the basic design of each intervention is detailed and intervention resources can be found at the DIVAS Program Resources website [28].

**Fig 1:**
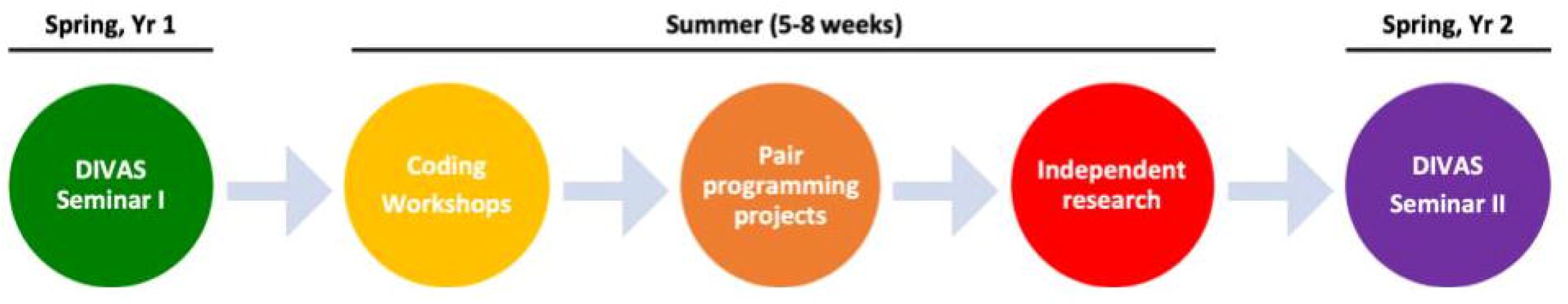
Interventions comprising the computational ‘onramp’ of the DIVAS program.

### DIVAS Seminar I and II

DIVAS Seminar I and II are both one-credit seminars offered in the spring semester. DIVAS Seminar I is offered to new scholars before the summer coding workshop and projects. DIVAS Seminar II is offered to scholars the spring after they have completed the summer interventions. DIVAS Seminar I is designed to introduce students to images as data and basic coding concepts, as well as allow them to meet professionals who use coding in their everyday work. Students complete a photo journal project where they identify a question or problem of interest, collect a series of images to address that question or problem, then use ImageJ to conduct simple image processing. In DIVAS Seminar II, students clean-up and annotate Python code written the previous summer. They also work with the instructor to make edits and improvements to the coding workshop as needed. Finally, students learn about and gain some familiarity with parallelization and grid computing. Course syllabi and sample resources for each course are available at the DIVAS Program Resources website [28].

### Coding Workshop

Short courses, such as those run by The Carpentries, have become a popular way to build coding and data analysis skills [29]. On average, participants report increased self-efficacy in coding and coding skills, based on pre- and post-workshop surveys and on longitudinal surveys [29,30]. However, workshops like those offered through The Carpentries are not targeted towards, nor significantly attended by, undergraduate students [29]. We designed a one-week coding workshop that includes two days of basic coding in Python and three days of image processing using OpenCV libraries. The two-day introduction to Python was modeled on an existing Carpentries workshop and can be found at GitHub [31]. The overall design of the three-day image processing workshop was informed by Adrian Rosebrook’s 2016 book on the topic [32]. To keep students engaged with Python basics, examples used during this section of the workshop were tailored toward image processing projects. Students were also presented with two authentic and “solvable” research problems at the beginning of the image processing portion of the workshop. For the first problem, participants were asked to count bacterial colonies on a plate image. For the second, participants were asked to track the progress of an acid-base titration captured on video. Our workshop design provides students an opportunity to immediately apply their recently acquired Python skills to write code to perform analysis tasks to address these two authentic problems. The image processing portion of the workshop was adopted by The Carpentries in 2019 [30,33]. At the same time, the image processing operations were translated into Scikit-image, which is much easier to install and implement across a wide range of hardware, software, and network environments. Workshop materials are available at its Data Carpentry site [30].

### Pair Programming Projects

Pair programming is a practice used in the software development industry in which two programmers work together, with one person assuming the role of the “driver” who writes the code, and the other taking the role of “observer” who reviews the code and makes suggestions. In introductory computer science courses, the use of pair programming results in higher quality code, increased student enjoyment, improved pass rates for courses, and improved retention in computer science majors for both men and women [17,34–36]. Also, pair programming has been shown to increase the confidence of women in the programming solutions they produce [34]. We designed the DIVAS program so that participants would transition from the coding workshop to pair programming work, applying knowledge gained in the workshop to the completion of two consecutive two-week pair programming projects. Each year, one project was morphometric in nature while the other was colorimetric. Image data sets were found from public repositories or from the research of the faculty team. The project was presented by a faculty member at the beginning of each project. DIVAS Scholars were randomly divided into pairs. For pairs composed of students at different institutions, pair programming was conducted virtually using Zoom. This arrangement allowed us to explore the feasibility of a fully online program. A significant amount of project management was done via the DIVAS Slack team. Each day, pairs met for a stand-up (brief 5-10 minute) meeting where progress and next steps were reported. Issues were also shared and discussed. Pairs worked on code for the remainder of the day. A formal code review was conducted each week by the DIVAS community of practice, with community members joining both in-person and virtually. All participants were to have copied and annotated the code of the other teams prior to the review. Progress and issues were discussed. As a group, major goals for the following week or final items to wrap up the project were identified. An example of a pair programming project can be found at the DIVAS Program Resources website [28].

### Independent Research

In year one, scholars were required to conduct 3-4 weeks of independent research after completing pair programming. Projects were based off of the existing research of the faculty team as well as were informed by student interest (Table 1). Students generally worked independently, but met with their faculty advisor for daily check-ins and to troubleshoot any problems that arose. Participating in DIVAS research was optional in years two and three to better accommodate student schedules, e.g. REU participation, study abroad, etc.

**Table 1.**
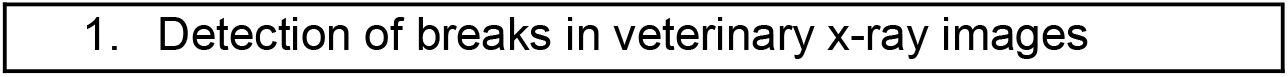
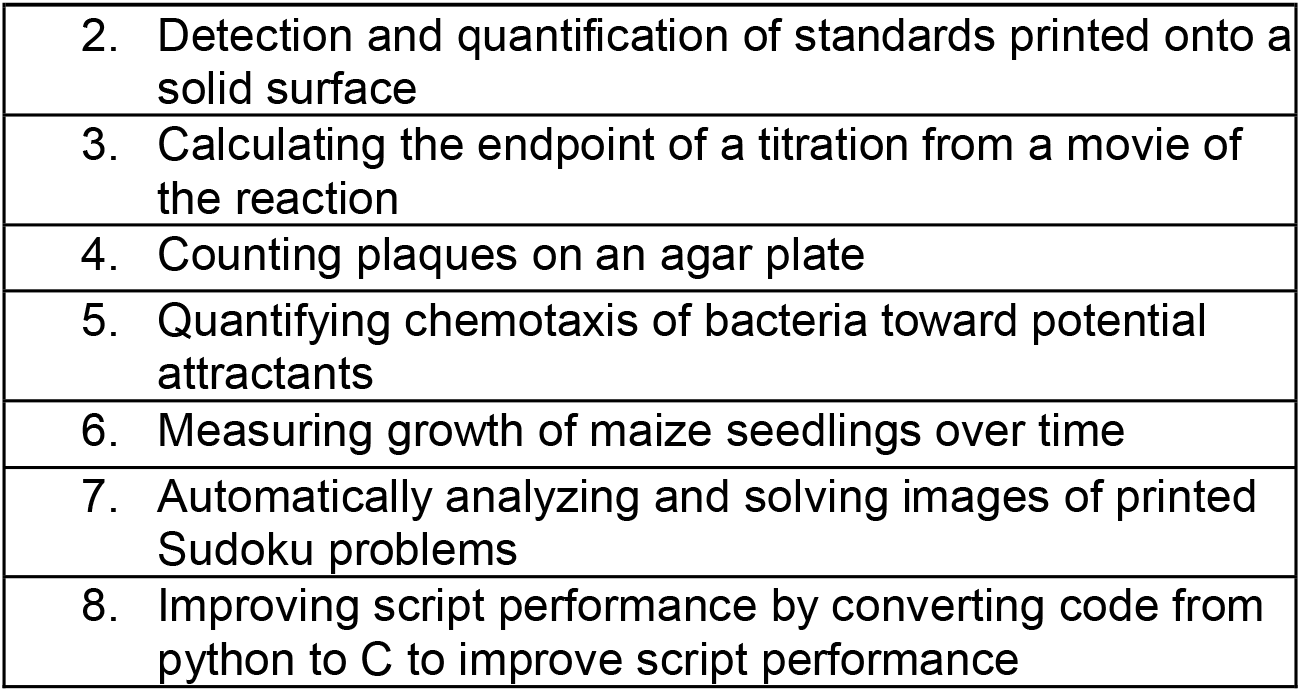
Example Pair Programming and Research Projects

|  |
| --- |
| 1. Detection of breaks in veterinary x-ray images |

|  |  |
| --- | --- |
| 2. | Detection and quantification of standards printed onto a solid surface |
| 3. | Calculating the endpoint of a titration from a movie of the reaction |
| 4. | Counting plaques on an agar plate |
| 5. | Quantifying chemotaxis of bacteria toward potential attractants |
| 6. | Measuring growth of maize seedlings over time |
| 7. | Automatically analyzing and solving images of printed Sudoku problems |
| 8. | Improving script performance by converting code from python to C to improve script performance |

Within the DIVAS Project framework, several questions were explored: 1) How do program interventions impact participant self-efficacy toward computation? 2) How do program interventions impact participant career interest? 3) How do program interventions and their ability to demonstrate effective computational thinking?

The overall objectives of the DIVAS project are to:

1. Explore the effectiveness of coding workshops on student attitudes toward computation and their ability to demonstrate effective computational thinking.
2. Measure impacts of paired programming projects, independent research, and professional development seminars on self-efficacy and ability to apply computational skills.
3. Investigate the impact of curricular and co-curricular interventions in computation on student preferred and actual career path.

## Materials and Methods

### Study Context

Each of the three years of the study, the DIVAS program was advertised using flyers (digital and paper), online and social media posts, and visits by faculty and existing scholars to classes that are generally enrolled by freshman and sophomore natural science majors. Up to six scholars were selected each year. Every effort was made to include each student who completed an application in the DIVAS program. A total of 17 scholars were selected to complete all program interventions (Table 2). Scholars were 76% women and 14% underrepresented minority (URM; African Americans, American Indians including Native Alaskans, Hispanics and Native Pacific Islanders). An additional 17 faculty, staff, and students participated in the one-week coding workshop and completed pre- and post-assessments.

**Table 2.**
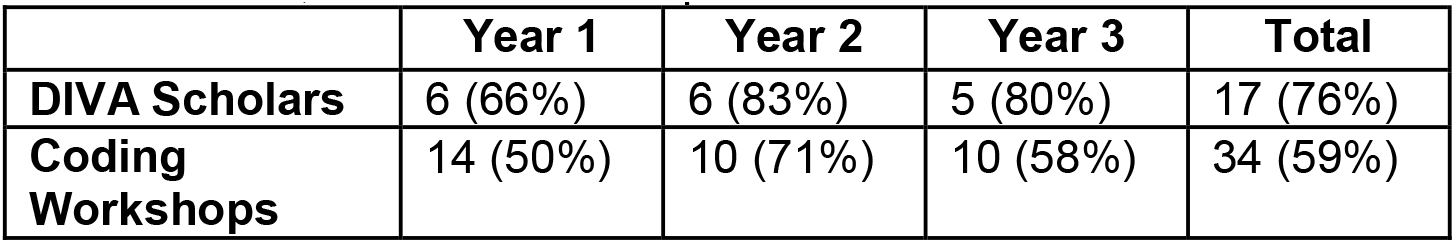
Participant overview. Participants who completed assessments, with % women in parenthesis.

|  | Year 1 | Year 2 | Year 3 | Total |
| --- | --- | --- | --- | --- |
| **DIVA Scholars** | 6 (66%) | 6 (83%) | 5 (80%) | 17 (76%) |
| **Coding Workshops** | 14 (50%) | 10 (71%) | 10 (58%) | 34 (59%) |

Scholars’ majors were biomedical engineering, biology, biochemistry, computer science, health and human performance, health and society, chemistry, and bioinformatics. STEM major retention was 100% at one year after the DIVAS II Seminar, with 89% of DIVAS Scholars retaining a STEM major for the duration of the program. Participants provided informed consent to provide self-efficacy and career path data by completing and submitting an electronic survey administered using Qualtrics software (Qualtrics, Provo, UT). Participants also provided informed consent to complete computational thinking prompts and to submit code generated for analysis, which was scored by project researchers. Doane University Institutional Review Board (IRB) approved the study.

### Self-efficacy and Career Path Assessment

A Qualtrics survey was used to measure perceived self-efficacy in computing and intention to pursue a career path involving computing. This survey, titled ‘DIVAS Career Path and Self-Efficacy’, is based on two previously-designed and validated surveys [37,38]. Survey questions ask participants to score their general knowledge of computational thinking and ability to use computational tools to solve problems; to indicate how much they know about careers using computer science applications, programming or computational thinking; and to assess how familiar they are with how to find information about computationally-related careers. Twelve questions related to self efficacy are answered as a user-inputted number on a 100-point scale, with higher values representing more self-efficacy for a particular item. Seven questions related to career paths include response choices on a four- or five-point Likert-type scale. Participants took the survey before and after the major interventions in the project. If the participant had previously completed the survey after an intervention, this score was used as the pre-survey for a subsequent intervention. A PDF of the survey can be found under ‘Assessment Tools’ at the DIVAS Program Resources website [28].

### Computational Thinking Assessment

A rubric was designed and iteratively revised to measure computational thinking based on definitions from the International Society for Technology in Education (ISTE) and Computer Science Teachers Association (CSTA), Carnegie Mellon, Google, and Harvard [39–42]. Our computational thinking (CT) rubric was organized into the first four phases of the RADIS (Recognize / Analyze / Design / Implement / Support) framework [43]. The Recognize section measures how well the problem is understood and one’s ability to gather the data needed to solve the problem. The Analyze section measures the ability of the participant to understand the options available to solve the given problem. This section also measures the ability of the participant to use abstraction, modeling/representation, and decomposition to design a solution to a problem. The Design section measures the participant’s ability to design an effective algorithmic procedure to solve the problem. It includes the participant’s ability to use sequence, selection, and iteration. The Implementation section addresses the ability of the participant to transform the algorithm into working code to solve a given problem. It also addresses the evidence that is used, reused, and remixed from previous projects or other sources. Finally, the Implement section assesses any testing or debugging that was used to improve the code. The original CT rubric was scored on a three-point scale; Proficient (3), Progressing (2), or Novice (1). Subsequent iterations included five levels, first from 0 to 4, then from 1 to 5. The additional levels were added to better accommodate the types of variation we were seeing in the scored artifacts. The expanded scale was adjusted to start with ‘1’ (indicating that something was attempted) from ‘0’ (indicating that nothing was attempted) to make statistical analysis more interpretable. Scales were standardized in the analysis so that group differences were comparable from year to year. The internal reliability of the first version and final versions of the instrument was high overall (Cronbach’s alpha of 0.94-0.95). Interrater reliability was determined to be 76% using a set of seven artifacts scored by three raters. The reliability of each section for both the first and final versions ranged from a Cronbach’s alpha of 0.80 for the ‘Implement’ section to a Chronbach’s alpha of 0.97 for the ‘Design’ section. The iterations of the CT rubric are available under ‘Assessment Tools’ at the DIVAS Program Resources website [28].

To assess CT ability before any formal instruction in coding, participants were given a handout that described a hypothetical cup stacking robot that could be given simple instructions to achieve different configurations of cups. The exercise was adapted from the Hour of Code lesson “Programming Unplugged: My Robotic Friends” [44]. Participants were asked to create a series of commands to achieve a particular cup stacking arrangement. After writing their initial set of commands, participants were asked to simplify their ‘code’, possibly by writing one or more new commands. A different cup-stacking prompt was used after the DIVAS Seminar I. After each subsequent intervention, the code developed in each one was used to assess CT ability. The cup stacking prompts are under ‘Assessment Tools’ at the DIVAS Program Resources website [28].

### Data Analysis

To investigate changes in student self-efficacy and career interest, scores within each category were summed to determine a composite score for each individual. A paired-samples t-test was performed (alpha = 0.05) to determine if composite scores before and after a given intervention were significant. For significant changes, the effect size was determined by calculating Cohen’s d. The change in CT scores within a year was determined by calculating a total score for an individual artifact from each rater and determining the median value. A paired-samples t-test was performed to determine if CT scored had changed after an intervention. Subscores within each of the areas of the rubric (Recognize, Analyze, Design, Implement) were also calculated and evaluated using a paired-samples t-test to determine whether significant changes within each area were observed. Effect sizes for significant differences were described by calculating Cohen’s d.

## Results and Discussion

Up to six scholars were selected each year of this three-year project, with a total of 17 scholars participating overall. Participant limits were due to budget constraints. Most scholars were women (76%) and 14% were a member of an URM. Most scholars (82%) were in their first year of college when starting the program. In addition to the DIVAS scholars, seventeen individuals participated in the coding workshop, 47% of whom were members of an URM group (Table 2). Scholars start the program with a one-credit seminar in the spring and end the program with a one-credit seminar the following spring, thereby participating in the full DIVAS pipeline as presented in Fig 1 (above).

The project measured the impact of DIVAS interventions on participant self-efficacy in using computation to solve problems, participants’ attitudes towards computation, and participant awareness/interest in computing careers using established instruments (see Materials and Methods). For all of the interventions of the DIVAS pipeline, self-efficacy toward computing and intended career path was assessed. Computational thinking was assessed in all of the interventions except DIVAS Seminar II. Pre- and post-assessments for each intervention were completed by participants. The post-test data from the prior intervention was used as a baseline for the next intervention in the pipeline. Overall, every intervention was found to have a positive effect on one or more measures (self efficacy, career interest, computational thinking) for at least one of the program years. In the proceeding sections, we discuss the assessment results for each intervention and conclude with overall program impacts.

### Self-efficacy and Career path data by intervention

#### DIVAS Seminar I

As described in DIVAS Program Elements, this seminar is the scholar’s entry point onto the DIVAS programmatic onramp. We saw significant gains in self-efficacy or career path each year of the program and in aggregate (Table 3).

**Table 3.** Self-efficacy and career path gains in DIVAS Seminar I. Effect sizes (Cohen’s-D) for each year of the program and the three years combined (Comb.) is shown. *, p < 0.5.

|  | Year 1 | Year 2 | Year 3 | Comb. |
| --- | --- | --- | --- | --- |
| **Self-efficacy** | 2.38* | ns | 0.74* | 0.35* |
| **Career path** | ns | 1.38* | ns | 0.63* |

An additional source of self-efficacy information came from the voluntary completion of a IDEA Student Ratings of Instruction system survey [45], which is conducted at Doane University at the end of each course and that we utilized in the DIVAS project. We analyzed self-reported learning gains in the IDEA-defined learning objectives for the eleven scholars who completed the survey (year 1 = 5, year 2 = 4, year 3 = 2). We found that scholars self reported strong gains in the objectives “Acquiring skills in working with others as a member of a team” and “Learning appropriate methods for collecting, analyzing, and interpreting numerical information.” The three cohorts rated both objectives at an average score of 4.45 out of 5 points. The Doane institutional average over the period of this project on these learning goals are 3.72 and 3.56, respectively. Overall, DIVAS Seminar I was effective in improving the self efficacy of Scholars toward computing, and positively influencing their intended career path. Observationally, the seminar was important in building rapport and a shared experience between all members (faculty and students) in the community of practice. In year three, the DIVAS cohort completed their photo diary project in tandem with 200-level graphic design students (Fig 2). This experience was one of several opportunities that the seminar provided to anchor image collection and analysis in relevant ways.

**Fig 2:**
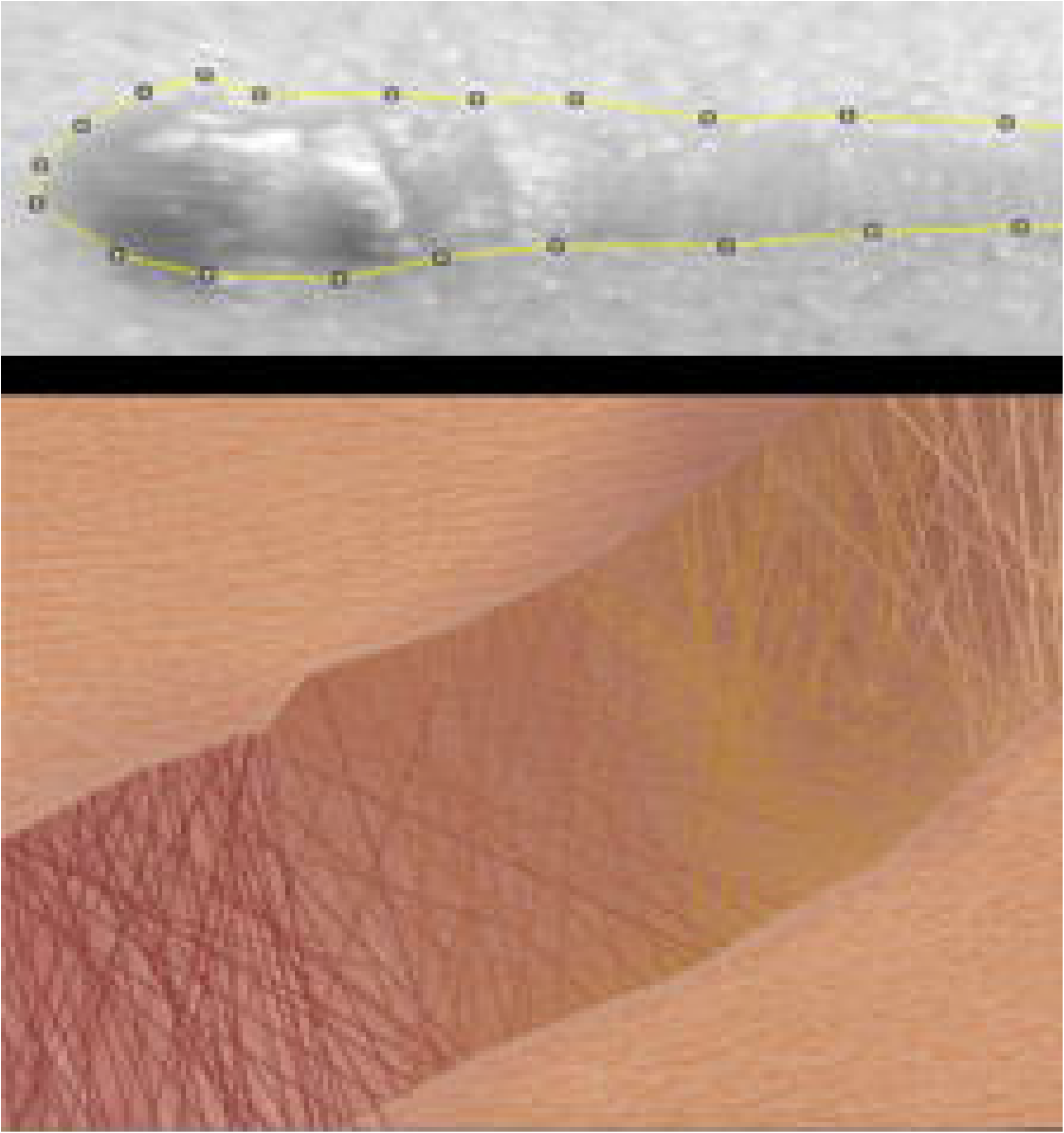
A collaborative photo journal project. A DIVAS scholar used ImageJ to analyze images of a healing wound (top) while a design student created a composition depicting the healing process (bottom).

#### Coding workshop

Modeled after existing Carpentries workshops, the five-day DIVAS workshop included two days of basic Python, Bash shell, and Git skills and a custom three-day workshop on basic computer vision topics using the OpenCV library for Python. The workshop was built around participants solving authentic research challenges within a community of practice focused on computation skills development (see ’DIVAS Program Elements’). We gathered self-efficacy and career path data pre- and post-workshop. We saw a significant improvement in self-efficacy in aggregate over the three years (Cohen’s-d = 0.57, p < 0.01). There were no significant changes in intended career path over the three year period. However, we did observe an increase in the standard deviation of the mean score. In looking at individual responses, this increase in standard deviation seems to indicate that scholars became more extreme at either end in their interest in incorporating computational skills in their future careers after this intervention. We did not find this concerning since this divergence in interest was paired with a significant increase in self-efficacy.

At the end of each day of the workshop, we also asked participants to rate the percentage of the day’s material they felt they had mastered. Data was compiled for all participants, including those who were not DIVAS scholars. We found a high average perceived mastery for the Python/Bash/git portion of the workshop, and then a drop for the first two days of the computer vision portion (Table 4). We believe this is due to the increased complexity in the subject matter. By the third day, this metric rose as participants were able to use their newfound skills to complete the challenge questions successfully.

**Table 4.** Participant responses to the question ‘What percentage of the day’s material do you feel you have mastered?’ for each day of the Coding Workshop.

|  | Day 1 | Day 2 | Day 3 |
| --- | --- | --- | --- |
| **Python Intro** | 75.9% | 76.3% | --- |
| **Image Processing** | 63.3% | 63.0% | 72.2% |

We found the coding workshop format to be effective as it immersed scholars in an enriching skill development environment. Though coding training was intensive, the participants’ self-reported improvements in mastery support the observation that scholars see tangible benefits from their persistence. The workshop also provided two cycles of challenge, learn, and achieve - in the spirit of Challenge Based Learning [46] - to provide participants multiple opportunities to struggle with new concepts and see the payoff.

#### Pair-Programming Projects and Research

Following the coding workshop, scholars employed pair programming to solve a colorimetric and a morphometric image analysis problem (Fig 3). For each problem, each pair developed code to extract relevant data from the images, analyze and present the data appropriately, and validate their results. To promote a community of practice, scholars participated in daily “stand-up” meetings where they gave brief progress reports and set goals for the day. Scholars participated in weekly guided code review sessions where they presented their code and provided critical feedback (written and verbal) on each pair’s code. The process was repeated with a second project with different partners in the following two weeks.

**Fig 3:**
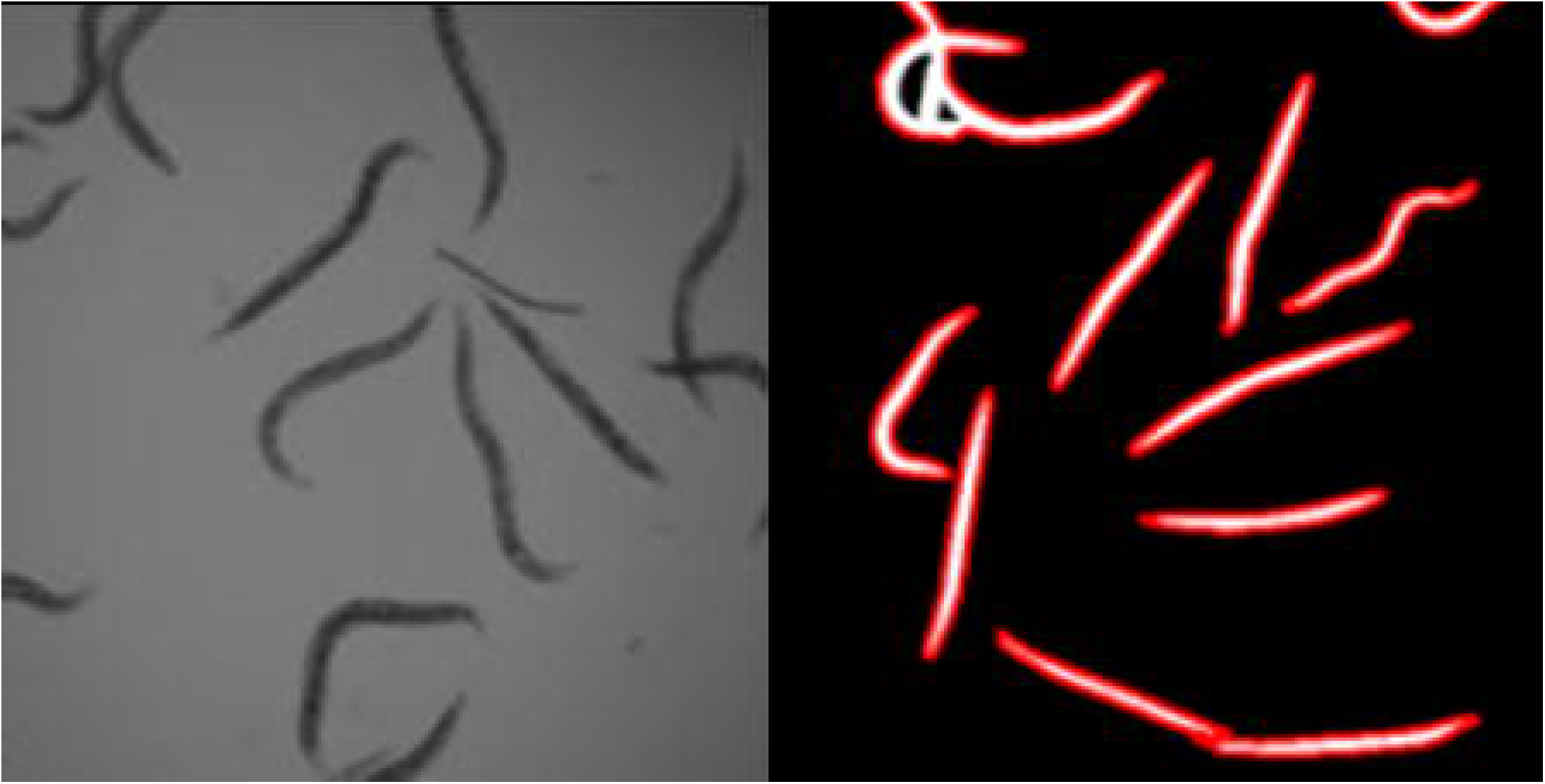
Example pair programming project in which students aimed to count the dead (straight) worms in a series of images. The unprocessed image is from [47].

Following pair programming work, scholars had the opportunity to complete three or more weeks of independent research (required in Year 1, optional in Years 2 and 3). Scholars either chose to work on existing projects, or design their own, within a faculty mentored research group. Examples of scholars’ projects included locating breaks in veterinary x-ray images, ingesting and solving printed Sudoku problems, greatly improving program performance by translating Python scripts into parallelized C++ code, and measuring chemotaxis of bacteria toward potential attractants (Table 1, above).

We collected self-efficacy and career path data upon the conclusion of summer research, if the scholar participated, or at the conclusion of pair programming projects for those students not participating in research. Although self-efficacy moved in a positive direction, we did not see significant gains in self-efficacy or career path at the end of summer activities. This was not surprising because students’ self-efficacy was already high and near the ceiling of the instrument, on average, following the coding workshop (Fig. 4). However, given that students were asked to solve a number of challenging problems largely independently, we see the maintenance of self-efficacy throughout this programmatic period as significant.

**Fig 4:**
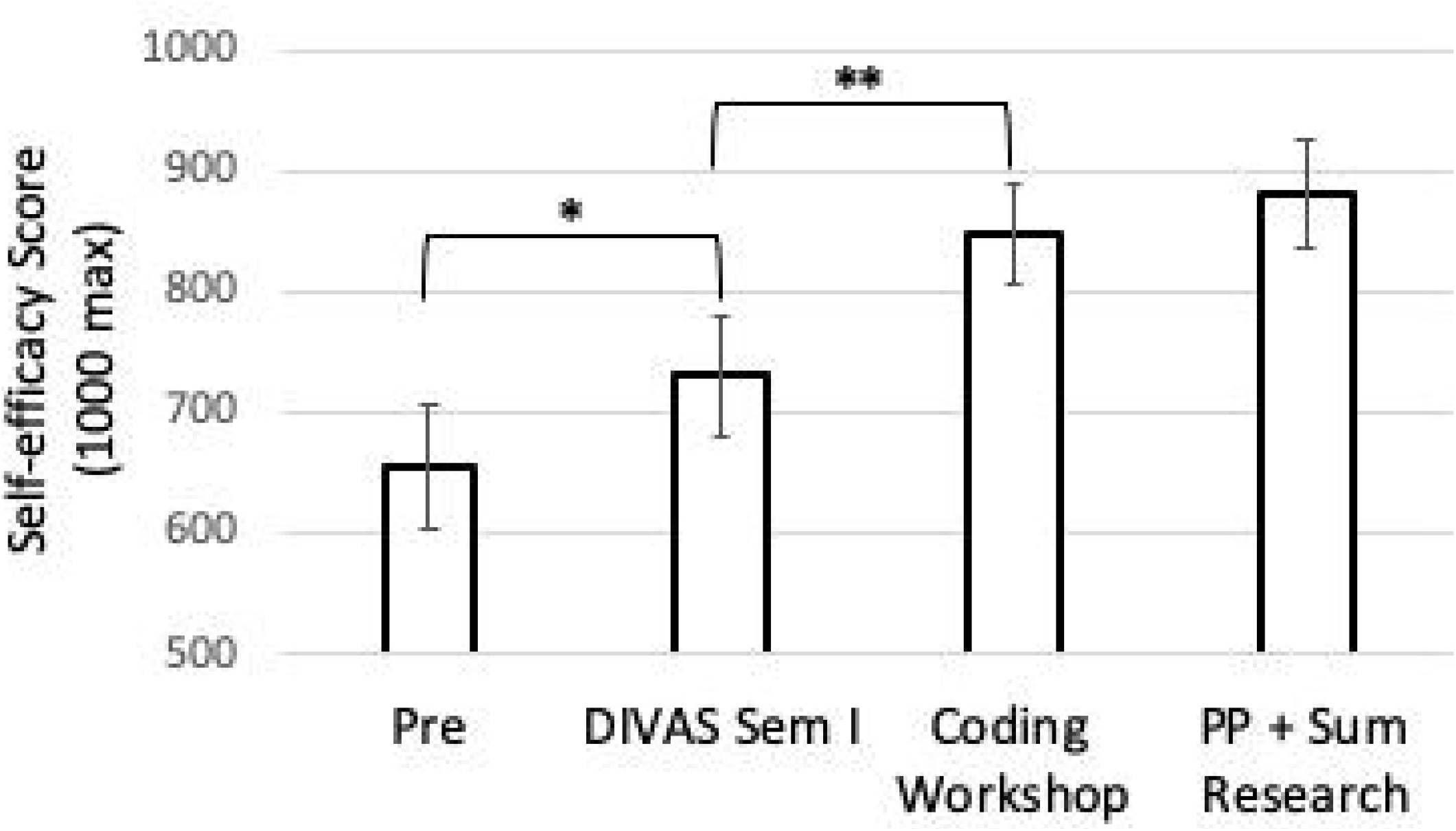
Average self-efficacy scores after DIVAS pipeline interventions for the three years of the project. Pre = pre-intervention score, PP = pair programming. *, p < 0.05; **, p < 0.01

Observationally, the pair programming and summer research projects were where scholars truly experienced the team-based environment of computational work. They learned to leverage each other’s ideas and expertise to develop approaches to solving a variety of problems. We found that scholars tended to work amongst themselves *before* seeking input from one of the faculty mentors. We considered this both a healthy development of independence and teamwork that reflected the increased confidence scholars gained in their individual and collective skill sets.

We also observed cases where one or more scholars would be given special authority by the group. While this was often productive, we also observed that it sometimes contributed to an over/underfunctioning dynamic between pairs. Because of this, we were especially mindful of giving praise for taking risks and highlighting the specific strengths of each project and each scholar separately. We also worked to minimize this over/underfunctioning dynamic when selecting pairs for each project so as to maximize each student’s engagement.

#### DIVAS Seminar II

In the first iteration of DIVAS Seminar II, scholars organized their project code repositories, developed online portfolios regarding their DIVAS experiences, and gave a local conference presentation. Based on student feedback, the next year’s Seminar II was modified to include more challenging academic content. Students learned parallel programming using Python and OpenMPI, creating a “Burning Ship” fractal image [48] using the Doane University supercomputer, Onyx. The third year, DIVAS Seminar II walked a line between the first two iterations; students worked on cleaning the previous summer’s code and keeping the code repository up to date, in addition to helping test the new version of the Image Processing workshop that used the Scikit-Image processing libraries versus OpenCV. DIVAS Seminar II was scheduled at the same time as DIVAS Seminar I each year. This made it easier to promote interactions between the classes. Further developing peer mentorship opportunities, the DIVAS faculty created a “writing center for computing” on campus called the Center for Computing in the Liberal Arts (CCLA)[49]. The center was led by a staff person hired, in part, to serve this role. Upon creation of the CCLA, several DIVAS Scholars signed up to serve as peer mentors, assisting in the creation of training materials and participating in center activities.

In addition to gains in self-efficacy and career path for the first cohort of scholars, a very significant gain in career path for second-year scholars was observed (Table 5). Similar to DIVAS Seminar I, responses on the IDEA survey for DIVAS Seminar II were also analyzed for perceived learning gains. Survey data showed that students perceived the largest gains in “Learning appropriate methods for collecting, analyzing, and interpreting numerical information” (4.33 ± 0.52). Scholars also responded positively to the statement, “My background prepared me well for this course’s requirements” (4.5 ± 0.55), reflecting the gains in self-efficacy we saw in the survey data. The last year of the seminar occurred during the first wave of the COVID-19 pandemic. This resulted in a response rate to the self-efficacy and career path survey that was too low to report.

**Table 5.** Self-efficacy and career path gains in DIVAS Seminar II. Effect sizes (Cohen’s-D) for Years 1 & 2. *, p < 0.5.

|  | Year 1 | Year 2 |
| --- | --- | --- |
| **Self-efficacy** | 1.2* | ns |
| **Career path** | 1.5* | 3.93** |

#### Computational thinking

CT ability was measured using a rubric designed by the team as described in the Materials and Methods. Participant responses were scored for CT ability for each intervention, except for DIVAS Seminar II, which did not include activities assessable via our rubric. In Year 1, we saw a significant improvement overall CT scores (Cohen’s d = 0.51, p < 0.1) and in the ‘implementation’ criteria for CT skills (Cohen’s d = 0.96, p<0.05) at the end of the coding workshop. We saw significant gains in overall CT scores after pair programming in both Years 2 (Cohen’s d = 2.0, p < 0.05) and 3 (Cohen’s d = 4.1, p < 0.05). There were significant gains in the ‘recognize’, ‘analyze’, and ‘design’ categories in year 2 (Cohen’s d = 1.5-2.1, p < 0.01). There were significant gains in the ‘analyze’, ‘design’, and ‘implement’ categories in year 3 (Cohen’s d = 2.44-4.57, p < 0.05).

#### Overall project outcomes and next steps

Over the three years of the project, scholars experienced significant increases in self-efficacy towards computing from the beginning of Seminar I to the end of summer programming (FIG 4). The most significant gains (p < 0.05) occurred during Seminar I and the coding workshops. The impact of the DIVAS program on scholars’ intended career paths was more subtle. Although scholars did not show significant career path gains from the initial pre-test before Seminar I to the end of summer research (p = 0.12), Seminar I resulted in significant gains in career interest for all years combined, as did Seminar II for Years 1 and 2 (Tables 3 and 5), both of which include explicit career exploration. Scholars were also observed to become ‘warmer’ or ‘colder’ to a career utilizing computing as they moved through the program. This effect is apparent in the increased standard deviation in post-intervention career interest scores, which started at ± 2.3 after Seminar I, grew to ± 3.02 after the coding workshop, and increased to ± 3.46 after pair programming/summer research. We see this as an encouraging progression, especially because scholar self-efficacy grew steadily throughout the program.

In a number of ways, DIVAS scholars have, persisted in coding and have incorporated skills gained in the DIVAS project into their academic careers, extracurricular activities, and career planning. One scholar majoring in biology declared a minor in software development. A second biology major switched to a bioinformatics major, and two scholars have taken non-required electives that emphasize computational skills. One scholar participated in an external REU program in computational and systems biology, and eight have continued research projects that incorporate coding or computational thinking. Three DIVAS scholars have worked as peer tutors for Doane’s CCLA. One former scholar is pursuing a Ph.D. in chemical biology with a significant computational component to their research and another student who participated in both the coding workshop and paired programming is pursuing a Ph.D. in complex biosystems.

Overall, even given the small sample represented in this study, we see great potential in the DIVAS approach of introducing novice students to computing through a media computing within a community of practice. Thirteen of the 17 DIVAS scholars from the three years of the project (76%) were women and 14% of scholars were members of an URM group, a significantly higher percentage than the total percentage of women and URM group members in the majors represented in the project or in the STEM workforce [50]. The large majority (82%) of scholars entered the program as freshmen. We retained 89% of DIVAS scholars for the duration of the program and retained 100% within a STEM major one year after completing the program. Our findings suggest that the DIVAS approach to computational skills development is a positive experience for students that warrants additional study through the implementation of DIVAS program elements in a broader array of educational contexts. To this end, we hope to form new DIVAS partnerships that will enable an expanded study on the efficacy of the DIVAS approach.

## Acknowledgements

The research described in this manuscript was funded by the National Science Foundation Digital Imaging and Vision Applications in Science Award IUSE-1608754. Students at Doane University participating in the project have also been supported by the National Science Foundation EPSCoR Center for Root and Rhizobiome Innovation Award OIA-1557417. The Welch Foundation (Grant# BH-0018) supported Dr. Burks and students of the Chemistry Department at St. Edward’s University. We would like to thank Adam Erck for his leadership in instituting the CCLA and Sarah Zulkoski for her tireless advocacy and support of this project.

## Supporting information captions

Not applicable.

